# VGLUT modulates sex differences in dopamine neuron vulnerability to age-related neurodegeneration

**DOI:** 10.1101/2020.11.11.379008

**Authors:** Silas A. Buck, Thomas Steinkellner, Despoina Aslanoglou, Michael Villeneuve, Sai H. Bhatte, Victoria C. Childers, Sophie A. Rubin, Briana R. De Miranda, Emma I. O’Leary, Elizabeth G. Neureiter, Keri J. Fogle, Michael J. Palladino, Ryan W. Logan, Jill R. Glausier, Kenneth N. Fish, David A. Lewis, J. Timothy Greenamyre, Antonello Bonci, Brian D. McCabe, Claire E. J. Cheetham, Thomas S. Hnasko, Zachary Freyberg

## Abstract

Age is the greatest risk factor for Parkinson’s disease (PD) which causes progressive loss of dopamine (DA) neurons, with males at greater risk than females. We found that vesicular glutamate transporter (VGLUT) expression mediates vulnerability to age-related DA neurodegeneration in a sex-dependent manner, providing a new mechanism for sex differences in selective DA neuron vulnerability. These findings lay the groundwork for novel therapeutic strategies to boost neuronal resilience throughout aging.

---

Progressive loss of midbrain dopamine (DA) neurons and their striatal projections are defining features of Parkinson’s disease (PD)^1^. Aging is the greatest risk factor for PD^2^, yet little is known about the mechanisms that determine DA neuron vulnerability across age. Additionally, not all DA neurons are equally susceptible to neurodegeneration^3-5^, as DA neurons in the ventral tegmental area (VTA) are more resilient than DA neurons of the substantia nigra *pars compacta* (SNc)^6,7^. Compared to SNc, a higher proportion of VTA DA neurons co-release glutamate and express vesicular glutamate transporter 2 (VGLUT2) which is responsible for the vesicular packaging of glutamate^8,9^. Notably, we previously showed that VGLUT2 also enhances vesicle loading and release of DA in response to increased neuronal activity^10,11^. However, the precise role of VGLUT2 expression in DA neuron resilience remains controversial. Adult DA neurons can upregulate VGLUT2 in response to cell stress as an adaptive, neuroprotective response to insult^12-14^. Nevertheless, other data suggest that excess VGLUT2 expression can induce DA neurodegeneration^13^. Moreover, while females have lower prevalence of PD, the molecular mechanisms underlying this sex difference remain unknown.

To answer these outstanding questions, we established an experimental system in *Drosophila* to dissect age-related DA neurodegeneration *in vivo*. As in mammals, DA neurons play a critical role in locomotion in *Drosophila*^15,16^, but it is unclear if flies also exhibit similar age-related and sex-specific motor differences. We therefore monitored 24-hour basal locomotion of male and female wild-type w^1118^ flies across the lifespan (Extended Data Fig. 1). We found a significant loss in basal locomotion as a function of aging and discovered that this age-related locomotor decline was greater in males than females (Extended Data Fig. 1). Consistent with mammals, these findings reveal greater susceptibility to age-dependent loss of locomotion in male *Drosophila*.

**Fig. 1.**
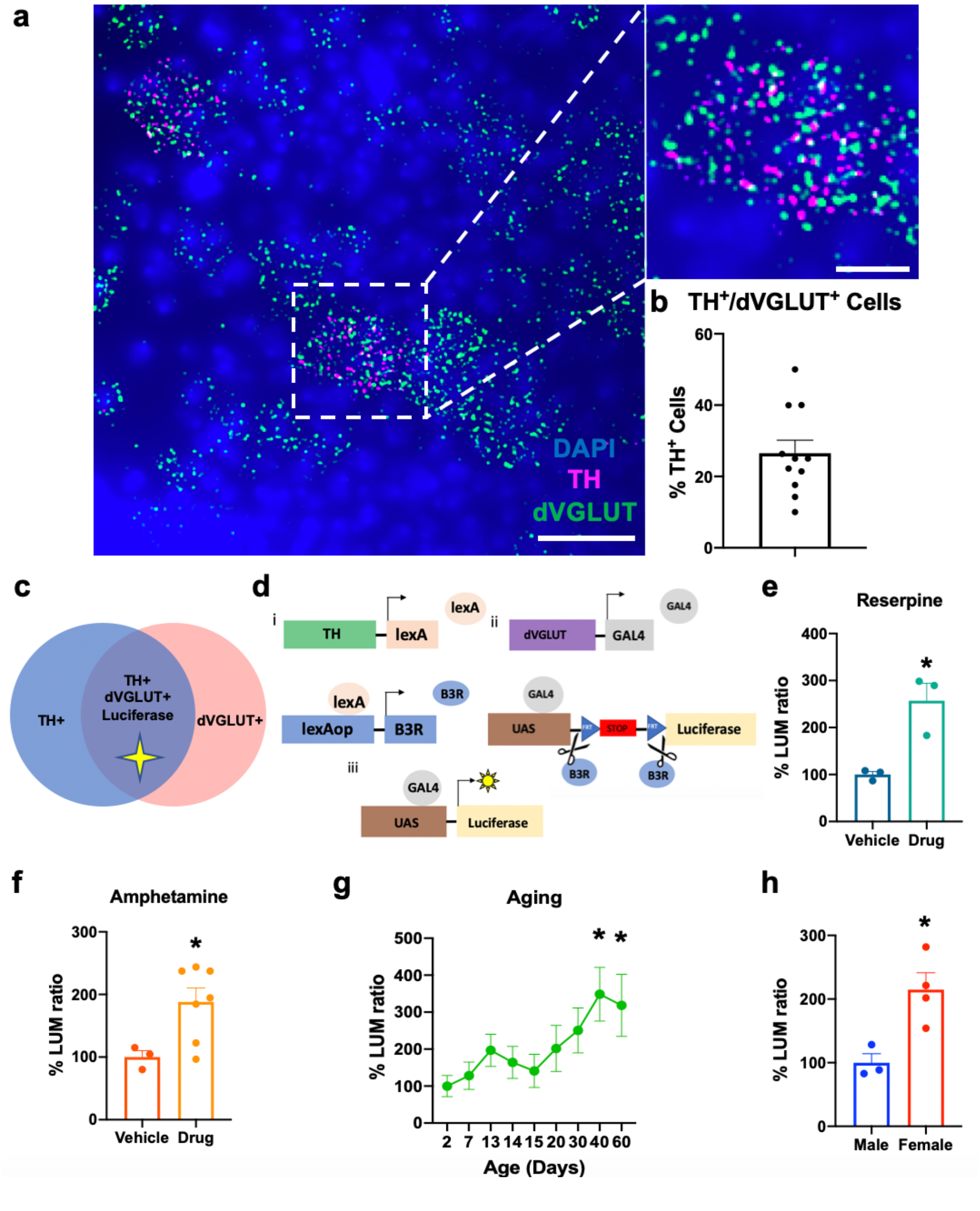
Intersectional genetic reporter shows altered DA neuron dVGLUT expression in response to vesicular DA depletion and to sex. **(a)** Representative confocal image demonstrating TH (magenta) and dVGLUT (green) mRNA expression via multiplex RNAscope in neurons of wild-type w^1118^ fly central brain 14 days post-eclosion; scale bar= 25 μm. Inset shows a zoomed-in cell expressing both TH and dVGLUT mRNA; scale bar=10 μm. **(b)** Quantification of the RNAscope labeling in adult central brain shows 27± 3.7% of TH^+^ DA neurons are also dVGLUT^+^. **(c)** Schematic where an intersectional genetic luciferase reporter of dVGLUT expression is expressed only in DA neurons that express both TH and dVGLUT. **(d) Panel i:** The TH promoter drives LexA to express B3 recombinase (B3R) in TH^+^ cells. **Panel ii:** B3R excises a transcriptional stop cassette within UAS-Luciferase. **Panel iii:** This permits successful dVGLUT-GAL4-driven transactivation of UAS-Luciferase selectively in TH^+^/dVGLUT^+^ cells. **(e)** Vesicular DA depletion by reserpine (300 μM, 24-hours) significantly upregulates dVGLUT expression in DA neurons compared to vehicle (2.6-fold increase; two-tailed unpaired t-test: t_4_=3.8, P=0.020). **(f)** Extended amphetamine treatment (10 mM, 24-hours) also increases DA neuron dVGLUT expression by 1.9-fold versus vehicle (two-tailed unpaired t-test: t_8_=1.9, P=0.049) **(g)** Increasing age progressively increases DA neuron dVGLUT expression (one-way ANOVA: F_8,49_=2.4, P=0.029) with a 3.4-fold increase in dVGLUT expression by day 40 versus day 2 post-eclosion (Bonferroni post-hoc test: P=0.023). **(h)** There are sex differences in DA neuron dVGLUT expression with female flies expressing 2.1-fold more dVGLUT compared to males at 14 days post-eclosion (two-tailed unpaired t-test: t_5_=3.4, P=0.019). Points represent individual animals. Results represented as mean± SEM; *P<0.05, **P<0.01; N=11 brains for RNAscope studies and N=3-7 samples per group in intersectional luciferase reporter studies.

We next investigated whether DA neuron-specific changes in VGLUT expression may be critical in age-related DA neurodegeneration in our *Drosophila* model. First, *in situ* hybridization via multiplex RNAscope confirmed the presence of tyrosine hydroxylase (TH)-expressing DA neurons that also express *Drosophila* VGLUT (dVGLUT) in the central brain of adult wild-type w^1118^ flies (Fig. 1a). Indeed, we identified 27± 3.7% of TH^+^ DA neurons are concurrently dVGLUT^+^ (Fig. 1b), in line with similar estimates of TH^+^/VGLUT2^+^ neurons in rodent midbrain^17,18^. We then employed a recombinase-induced intersectional genetic method to express a reporter of dVGLUT expression specifically in TH^+^/dVGLUT^+^ DA neurons (Fig. 1c). Since TH is the rate-limiting enzyme of DA biosynthesis, we constructed a fly strain using the TH-LexA expression driver, where B3 recombinase (B3R) is targeted to TH^+^ DA neurons to excise a stop cassette. This permits expression of a firefly luciferase transcriptional reporter driven by the dVGLUT promoter only in TH^+^ DA neurons (Fig. 1d). Consequently, our intersectional luciferase reporter produces a robust DA neuron-specific luminescent indicator of dVGLUT expression level with high signal-to-noise when compared to controls (Extended Data Fig. 2). Expression of the dVGLUT reporter is consistent with previous work in *Drosophila* demonstrating dVGLUT protein in a subset of DAergic nerve terminals^10^.

**Fig. 2.**
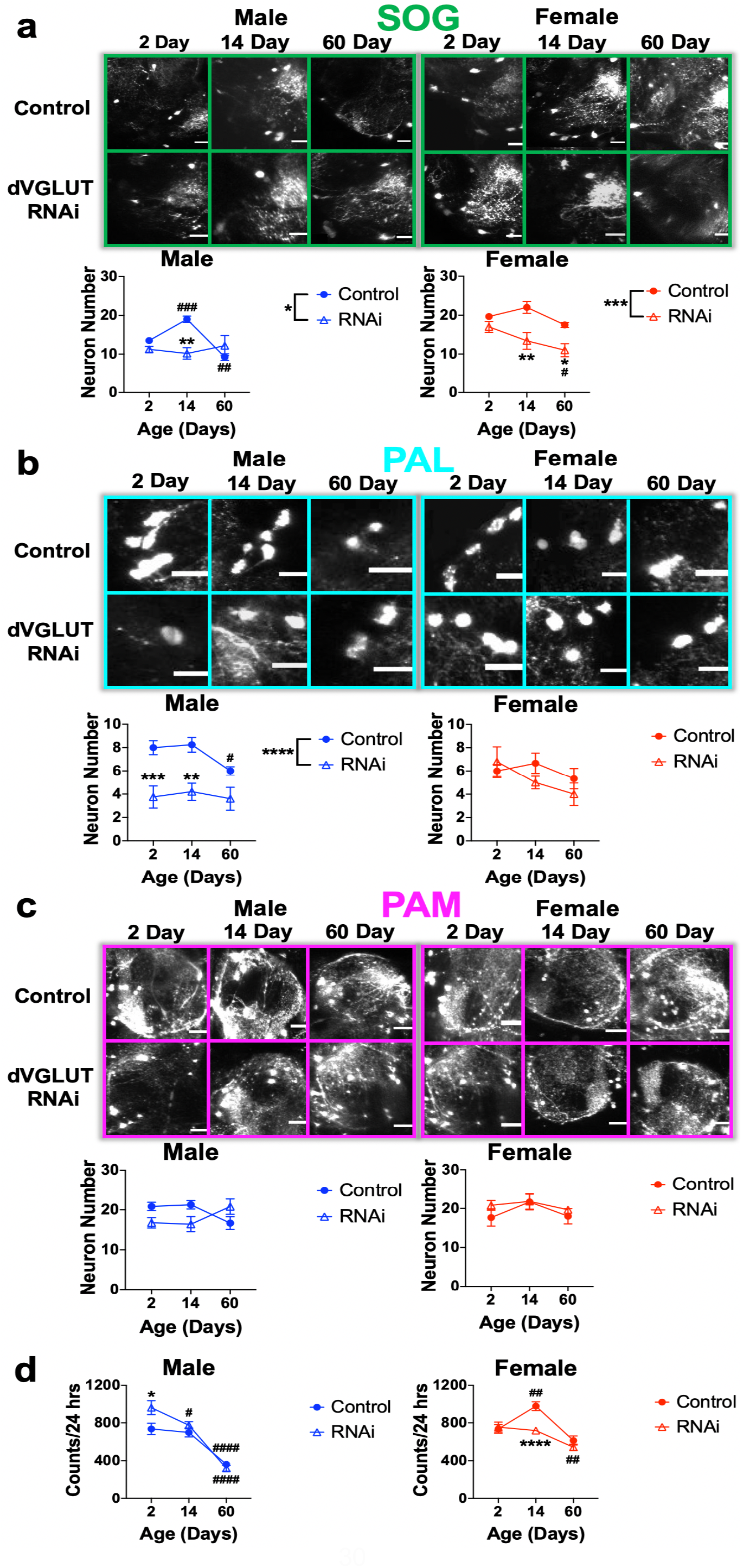
dVGLUT RNAi knockdown diminishes sex- and region-selective DA neuron resilience in aging. **(a)** Representative multiphoton images of GFP-labeled DA neurons within the SOG of whole *Drosophila* adult central brain from flies with the following genotypes: TH-GAL4/UAS-GFP (Control) (**Top Row**) versus TH-GAL4,UAS-GFP/UAS-dVGLUT RNAi (dVGLUT RNAi) (**Bottom Row**); scale bars=20 μm. Three-way ANOVA: Age: F_2,42_=6.0, P=0.0049; Sex: F_1,42_=20.9, P<0.0001; dVGLUT RNAi: F_1,42_=22.0, P<0.0001; Age×dVGLUT RNAi: F_2,42_=5.6, P=0.0068; Age×Sex: F_2,42_=0.9, P=0.40; Sex×dVGLUT RNAi: F_1,42_=3.1, P=0.087; Age×Sex×dVGLUT RNAi: F_2,42_=3.0, P=0.061. **(b)** Representative multiphoton images of PAL DA neurons; scale bars=20 μm. Three-way ANOVA: Age: F_2,45_=3.3, P=0.044; Sex: F_1,45_=0.0, P=0.99; dVGLUT RNAi: F_1,45_=18.4, P<0.0001; Age×dVGLUT RNAi: F_2,45_=0.5, P=0.62; Age×Sex: F_2,45_=0.3, P=0.75; Sex×dVGLUT RNAi: F_1,45_=8.0, P=0.0070; Age×Sex×dVGLUT RNAi: F_2,45_=1.4, P=0.26. **(c)** Representative multiphoton images of PAM DA neurons; scale bars=20 μm. Three-way ANOVA: Age: F_2,45_=0.9, P=0.39; Sex: F_1,45_=1.5, P=0.23; dVGLUT RNAi: F_1,45_=0.0, P=0.98; Age×RNAi: F_2,45_=02.8, P=0.073; Age×Sex: F_2,45_=0.9, P=0.43; Sex×dVGLUT RNAi: F_1,45_=3.0, P=0.093; Age×Sex×dVGLUT RNAi: F_2,45_=2.6, P=0.088. **(d)** Locomotion over a 24-hour period in male and female control and dVGLUT RNAi flies. Three-way ANOVA: Age: F_2,427_=81.1, P<0.0001; Sex: F_1,427_=9.7, P=0.002; dVGLUT RNAi: F_1,427_=0.1, P=0.75; Age×dVGLUT RNAi: F_2,427_=5.3, P=0.0052; Age×Sex: F_2,427_=12.4, P<0.0001; Sex×dVGLUT RNAi: F_1,427_=13.0, P=0.00034; Age×Sex×RNAi: F_2,427_=3.5, P=0.031. Results represented as mean± SEM; *P<0.05, **P<0.01, ***P<0.001, ****P<0.0001 compared to control by Bonferroni post-hoc test, ^#^P<0.05, ^##^P<0.01, ^####^P<0.0001 compared to day 2 post-eclosion by Bonferroni post-hoc test. N=3-8 brains per group for neuron counts and N=24-49 flies for locomotion.

Using our luciferase reporter, we first examined whether DA neurons dynamically alter dVGLUT expression following depletion of vesicular stores of DA. Treatment with reserpine (300 μM, 24 hours), which irreversibly blocks VMAT to prevent vesicular DA loading^19^, leads to a 2.6-fold increase in reporter output compared to vehicle (Fig. 1e). Similarly, amphetamine (10 mM, 24 hours), which depletes presynaptic DA stores over time by disrupting the vesicular pH gradient (*Δ*pH)^19^, causes a 1.9-fold increase in luciferase reporter expression (Fig. 1f). Next, in the context of aging, we observed a 3-fold increase in luciferase reporter expression in 40- and 60-day post-eclosion adult flies compared to young flies (day 2 post-eclosion) (Fig. 1g), suggesting a maximum level of dVGLUT upregulation is reached in DA neurons by 40 days. Together, our findings demonstrate that dVGLUT expression in DA neurons is dynamic. Importantly, these data also raise the possibility that DA neuron dVGLUT expression is elevated as a compensatory response to progressive loss of synaptic DA due to age-related DA neurodegeneration.

Since female flies show diminished vulnerability to age-related locomotor decline relative to males, and DA neurons upregulate dVGLUT during aging, we examined whether there are sex differences in DA neuron dVGLUT expression. Indeed, we discovered that adult females express 2.1-fold more dVGLUT in DA neurons compared to males at day 14 post-eclosion (Fig. 1h). We also asked whether these differences are conserved across species. We found that in human VTA/SNc, the relative density of TH^+^/VGLUT2^+^ neurons is 6.6 times greater in females than in males (Extended Data Fig. 3a, b). Furthermore, adult female rats similarly express 2.4-fold more VGLUT2 protein versus age-matched males in SNc TH^+^/VGLUT2^+^ DA neurons (Extended Data Fig. 3c), demonstrating evolutionarily conserved sex differences in DA neuron VGLUT expression.

**Fig. 3.**
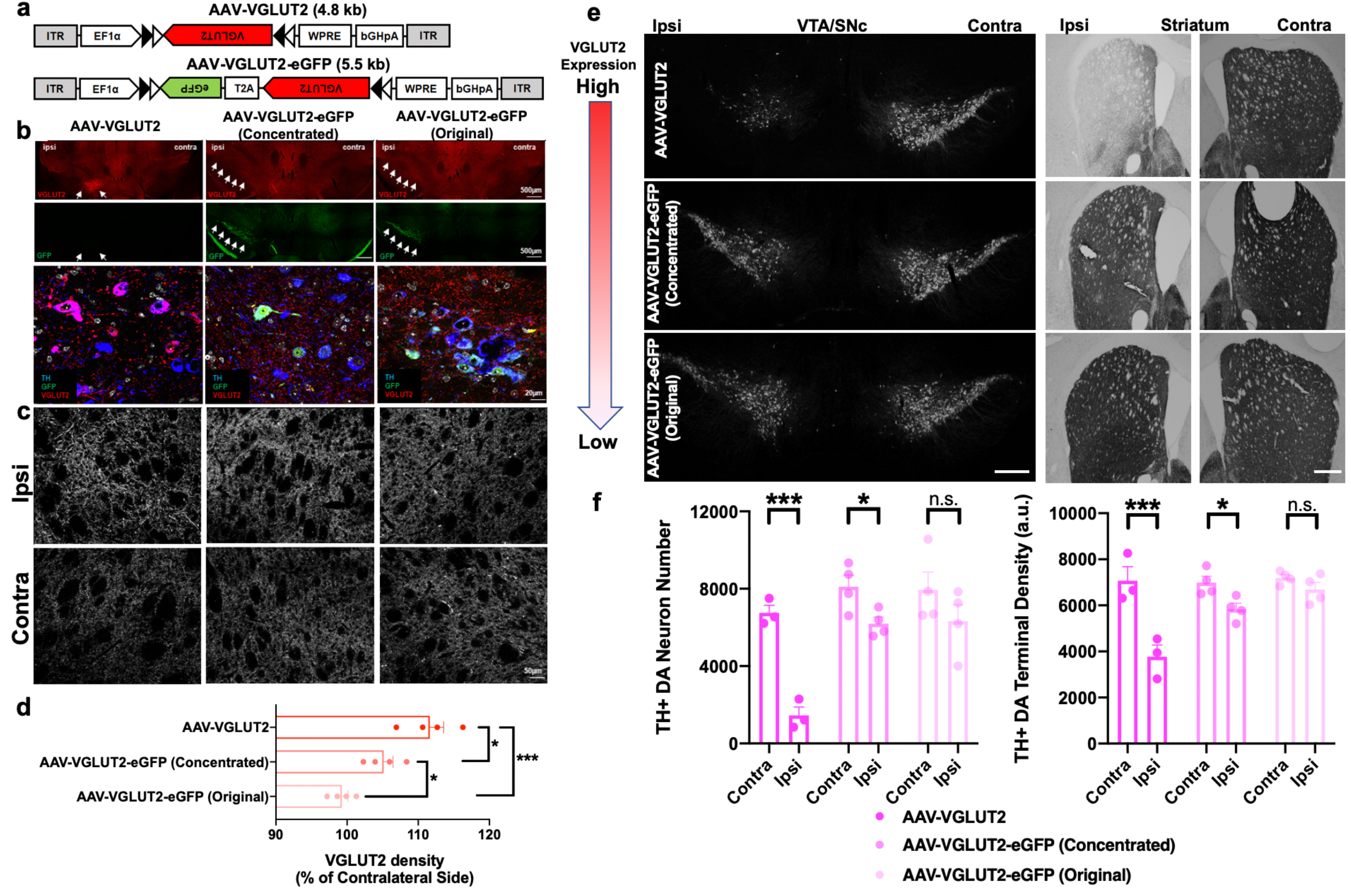
DA neuron survival in response to different levels of AAV-mediated VGLUT2 overexpression. **(a)** Schematic of AAVs used for cell-type specific VGLUT2 overexpression in SNc DA neurons of DAT^Cre^ mice. **(b, c)** Detection of VGLUT2 protein in **(b)** cell bodies and **(c)** striatal terminals of SNc DA neurons after unilateral injection of AAV-VGLUT2 (**left panel**), AAV-VGLUT2-eGFP concentrated titer (**middle panel**), and AAV-VGLUT2-eGFP original titer (**right panel**) in DAT^Cre^ mice. Ipsi indicates ipsilateral site of viral injection; contralateral side is the uninjected control. **(d)** Quantification of striatal VGLUT2 density relative to uninjected contralateral sites. One-way ANOVA: F_2,9_=18.2, P=0.0007. **(e)** Representative midbrain (**left panel**) and striatal sections (**right panel**) labeled for TH in mice unilaterally injected with either AAV-VGLUT2 (upper panel), concentrated titer AAV-VGLUT2-eGFP (middle panel) or original titer AAV-VGLUT2-eGFP (lower panel); scale bars=500 μm. **(f) Left panel:** Quantification of TH^+^ neuron number (left panel). Two-way ANOVA: Vector: F_2,8_=7.5, P=0.015; Hemisphere: F_1,8_=76.2, P<0.0001; Vector×Hemisphere: F_2,8_=11.2, P=0.0048. **Right panel:** Quantification of striatal density of TH^+^ DA nerve terminals. Two-way ANOVA: Vector: F_2,8_=7.1, P=0.017; Hemisphere: F_2,8_=51.4, P<0.0001; Vector×Hemisphere: F_2,8_=12.4, P=0.0036. Points represent individual animals with N=3-4 animals per group. Results represented as mean± SEM. *P<0.05, ***P<0.001 compared by Bonferroni post-hoc test.

We next determined whether there are also sex differences in DA neuron survival across aging, and if loss of dVGLUT expression in DA neurons diminishes such sex differences. For these experiments, the TH promoter drove expression of a GFP marker to selectively label DA neurons in whole living brain preparations from adult flies aged 2, 14 and 60 days post-eclosion (Extended Data Fig. 4). We examined several well-defined DA neuron clusters within the adult central fly brain associated with classic DAergic functions including locomotion and reward processing: the suboesophageal ganglion (SOG) cluster, protocerebral anterior lateral (PAL) cluster, and the protocerebral anterior medial (PAM) cluster. Using this experimental system, we next knocked down dVGLUT expression specifically in DA neurons via previously validated dVGLUT RNA interference (RNAi)^10^. We combined UAS-GFP and the TH-GAL4 driver strains via genetic recombination and crossed the progeny to dVGLUT RNAi flies, enabling us to image GFP-labeled DA neurons where dVGLUT expression has been knocked down. As a control, we confirmed that recombining TH-GAL4 with GFP does not alter DA neuron numbers across brain regions compared to non-recombined flies (termed ‘control flies’) (Extended Data Fig. 5).

We discovered that dVGLUT knockdown in DA neurons renders SOG DA neurons more vulnerable to age-related neurodegeneration (Fig. 2a). Interestingly, males show an increase in GFP-labeled DA neurons at 14 days compared to 2 days post-eclosion (P=0.0002), consistent with ongoing DA neuron development after eclosion^20^. By day 60, however, males have fewer SOG DA neurons compared to day 2 (P=0.0028). Moreover, DA neuron dVGLUT knockdown renders male SOG DA neurons more vulnerable to age-related DA neurodegeneration. Male dVGLUT RNAi flies demonstrate 50% fewer SOG DA neurons at day 14 post-eclosion compared to control males (P=0.0011). Critically, whereas control females show no significant DA neuron loss in the SOG across aging (P>0.05), dVGLUT RNAi females exhibit progressive DA neurodegeneration with 35% of SOG DA neurons lost by day 60 post-eclosion (P=0.030). In addition, dVGLUT RNAi females have significantly fewer SOG DA neurons compared to controls at 14 days (P=0.0089) and 60 days (P=0.027) post-eclosion. In the PAL, similar to the SOG, control males show a significant age-related decrease in PAL DA neurons at day 60 post-eclosion compared to day 2 (P=0.042), while females are protected with no DA neurodegeneration over time (P>0.05) (Fig. 2b). In males, dVGLUT knockdown in DA neurons leads to consistent reduction of PAL DA neurons across all timepoints compared to male controls (P<0.0001), suggesting a role for dVGLUT in male DA neuron development. In contrast, no significant differences in PAL DA neurons were observed between female dVGLUT RNAi versus control flies (P>0.05). In the PAM, we found no significant effects in males and females (P>0.05) (Fig. 2c), indicating that PAM DA neurons are not significantly impacted by age or sex.

Lastly, consistent with our imaging data, dVGLUT RNAi-mediated knockdown in DA neurons attenuates females’ apparent protection from age-related declines in locomotor behavior (Fig. 2d). dVGLUT RNAi females exhibit significant declines in locomotion at day 60 post-eclosion compared to day 2 (P=0.0070), whereas control females do not show significant age-related changes (P>0.05). In contrast to females, control male dVGLUT RNAi flies demonstrate significant locomotor decreases with aging. However, DA neuron dVGLUT knockdown accelerated these age-related locomotor declines with diminished locomotion observed at both days 14 (P=0.017) and 60 (P<0.0001) in dVGLUT RNAi flies compared to only day 60 (P<0.0001) in controls relative to day 2 post-eclosion. In all, these results suggest that adult *Drosophila* show clear region- and sex-specific differences in age-related DA neuron loss, and our results point to dVGLUT’s importance in both sex differences and age-mediated increases in DA neuron vulnerability.

Our data agree with findings in mice showing that VGLUT2-expressing DA neurons are more resilient to insults^12-14^. We previously examined whether boosting VGLUT2 expression in the SNc via heterologous overexpression could further increase DA neuron resilience. However, while Shen *et al*. showed that VGLUT2 overexpression increases DA neuron resilience to DA neurotoxin 1-methyl-4-phenyl-1,2,3,6-tetrahydropyridine (MPTP)^14^, Steinkellner *et al*. found that VGLUT2 overexpression selectively increases DA neuron vulnerability^13^. In determining why these two results are at odds with one another and our current findings concerning VGLUT’s role in DA neuron vulnerability, a key clue may lie with the respective viruses used in these earlier studies. Indeed, the two studies relied on viruses of different genomic sizes and titers where Steinkellner *et al*. employed AAV-VGLUT2 (genome size 4.8kb, 3.3×10^12^ gc/ml)^13^, while Shen and colleagues used AAV-VGLUT2-eGFP (genome size 5.5kb, 2×10^13^ gc/ml)^14^ (Fig. 3a). To determine if the viral preparations have conflicting effects because they produce differing levels of VGLUT2 overexpression, we unilaterally injected these Cre-dependent viruses into the SNc of mice expressing Cre recombinase under the control of the dopamine transporter (DAT^Cre^) for DA neuron-selective VGLUT2 overexpression.

Examination of VGLUT2 protein following viral transduction revealed a virus-dependent gradient of VGLUT2 overexpression in DA neurons and their terminals on the ipsilateral injected side. Highest levels of VGLUT2 overexpression were generated by AAV-VGLUT2 despite a lower titer (3.3×10^12^ gc/ml) compared to AAV-VGLUT2-eGFP (2×10^13^ gc/ml). Intermediate VGLUT2 overexpression was produced by a titer of AAV-VGLUT2-eGFP that was 2.5-fold more concentrated than originally used by the Shen *et al*. study (5×10^13^ gc/ml), with the least overexpression found with the original titer of AAV-VGLUT2-eGFP (2×10^13^ gc/ml)(Fig. 3b,c). While it is important to note that this approach doesn’t discriminate between endogenous (*e*.*g*., from thalamic and midbrain inputs) and heterologously expressed VGLUT2, we nevertheless detected a main effect of AAV-VGLUT2 vector type (Fig. 3d), suggesting we achieved different levels of DA neuron VGLUT2 overexpression *in vivo* using different AAVs.

We investigated whether the different levels of VGLUT2 overexpression produced by the AAVs impact DA neuron survival. Transduction with AAV-VGLUT2 led to a 79± 6.4% decrease in SNc DA neuron number (P=0.0001) (Fig. 3e,f), similar to our previous report^13^. In contrast, the original AAV-VGLUT2-eGFP did not induce significant DA neuron loss compared to the uninjected contralateral side (P>0.05), also consistent with earlier reported results^14^. We discovered that intermediate levels of VGLUT2 overexpression via the concentrated AAV-VGLUT2-eGFP virus reduced TH^+^ DA neuron number by 24± 4.1% (P=0.026). Consistent with the respective AAVs’ effects on DA neuron number, there was a 47± 7.2% loss of TH^+^ DA nerve terminal density in the striatum after AAV-VGLUT2 vector administration (P=0.00020), compared to 16± 3.7% loss for the concentrated AAV-VGLUT2-eGFP (P=0.049). Crucially, we observed no significant changes in either DA neuron numbers or terminal density for the original AAV-VGLUT2-eGFP titer (P>0.05) (Fig. 3e,f). Our data therefore suggest that DA neurons require a finely tuned balance of VGLUT2 expression to effectively boost resilience.

Together, our findings demonstrate sex differences in DA neuron vulnerability to age-related neurodegeneration. We provide evidence that VGLUT expression is a critical factor for the greater resilience of females to DA cell loss and associated locomotor impairments. Furthermore, we show sex differences in DA neuron VGLUT expression where DA neurons of female flies, rats, and humans express significantly higher levels of VGLUT compared to males. Finally, we demonstrate that the delicate balance of VGLUT2 expression in DA neurons is critical for its effects on resilience. This suggests a “Goldilocks” model: too little VGLUT upregulation may be insufficient to boost neuronal resilience to successfully weather periods of cell stress. When VGLUT2 expression is optimally enhanced in a physiological range, this may compensate for DA system degeneration and enhance DA neuron resilience, as supported by a recent study^12^. On the other hand, either too much VGLUT expression or prolonged duration of overexpression may leave the cells more vulnerable. Targeting this VGLUT-dependent mechanism therefore provides opportunities for new therapeutic approaches to boost neuronal resilience in both healthy aging and PD, while also revealing fundamental insights into the unique properties of DA/glutamate neurons.

## Methods

### *Drosophila* Experiments

#### Drosophila strains

All *Drosophila melanogaster* strains were grown and maintained on standard cornmeal-molasses media at 24°C, ~50% humidity under a 12:12 hour light/dark cycle in humidity- and temperature-controlled incubators. Unless otherwise noted, fly stocks were obtained from the Bloomington Stock Center. We used the wild-type w^1118^ strain for locomotor behavioral assays. We also used the following previously described transgenic stocks: *TH-GAL4* (gift of Dr. S. Birman, Université Aix-Marseille II-III, Marseille, France)^21^ and *TH-LexA*^22^ to drive expression in DA neurons via the GAL4/UAS and LexA/LexAop binary expression systems, respectively. *dVGlut-GAL4* was used to drive expression in dVGLUT-expressing neurons^23,24^. For imaging experiments, to label DA neurons with GFP, *UAS-GFP* ^25^ was genetically recombined with the *TH-GAL4* expression driver on chromosome III to construct the *TH-GAL4,UAS-GFP* fly strain. To ascertain effects of dVGLUT RNAi knockdown on age-related vulnerability of GFP-labeled DA neurons, we crossed *TH-GAL4,UAS-GFP* and *UAS-VGLUT-RNAi* (*UAS-Vglut-RNAi*^*HMS*^, HMS02011, VALIUM20, target 3077-3098 nt, chromosome 3)^10,26^ strains to generate *TH-GAL4,UAS-GFP/UAS-VGLUT-RNAi* flies; as a control, *TH-GAL4,UAS-GFP* flies were crossed to wild-type w^1118^ (*TH-GAL4,UAS-GFP/*+). All fly strains were outcrossed for 10 generations into the w^1118^ wild-type genetic background.

#### Construction of transgenic Drosophila strains

*Construction of LexOP>B3R strain*. B3 recombinase was amplified from pJFRC157-20XUAS-IVS-B3::PEST^27^ (Addgene plasmid #32136) using primers to add a syn21 translational enhancer sequence and remove the PEST domain. The resulting PCR product was transferred into pBID1-13xLexOP, a derivative of the pBID expression vector^28^ containing 13 LexOP sequences. This construct was introduced into the attP40 site on chromosome II by phiC31 injection (Genetivision, Houston, TX).

##### Construction of UAS>B3RT-STOP-B3RT-Luciferase strain

The *UAS>B3RT-STOP-B3RT-Luciferase* strain was created by replacement of the myr::RFP sequence in pJFRC160 (Addgene plasmid #32139)^27^ with firefly luciferase. This modified construct was introduced into the attP2 site on chromosome III by phiC31 injection (Genetivision).

##### Construction of the DA neuron dVGLUT luciferase reporter strain

To measure changes in DA neuron dVGLUT expression, we assembled our intersectional genetic reporter of DA neuron dVGLUT expression using a luciferase reporter: *dVGLUT-GAL4/LexOP>B3R;TH-LexA/UAS>B3RT-STOP-B3RT-Luciferase*. The B3 recombinase recognizes sequence-specific recombination target sites, B3RTs, in a highly specific manner and excises the intervening DNA between recombination sites, leaving behind a single recombinase target site^27^. Since TH-LexA-driven B3 recombinase is only expressed in DA neurons, this results in cell-specific excision of the B3RT-flanked STOP cassette, enabling dVGLUT-GAL4-driven expression of the intersectional luciferase reporter only in DA neurons.

#### Drosophila behavior

To monitor the locomotor response to aging, we used the Trikinetics *Drosophila* Activity Monitoring (DAM) system (Trikinetics, Waltham, MA) as described earlier^10^. Newly eclosed male and female flies were collected daily at fixed, regular times to ensure precise age determination. Adult flies were entrained in 12:12 hour light:dark (LD) cycles at 24°C, ~50% humidity in a humidity- and temperature-controlled incubator. For experiments employing the wild-type w^1118^ strain, flies aged 2, 14, 30 or 60 days post-eclosion were transferred individually into activity tubes containing standard cornmeal-molasses media which were then placed into activity monitors (DAM5, Trikinetics); studies examining locomotor behavior of DA neuron-specific dVGLUT RNAi (*TH-GAL4,UAS-GFP/UAS-dVGLUT-RNAi*) and control (*TH-GAL4/UAS-GFP*) were collected at 2, 14 or 60 days post-eclosion. Flies were allowed to acclimate to the monitors and individual recording vials for 24-hours followed by continuous 24-hour activity monitoring (also in LD) for movement. Locomotor activity was measured as the number of times (counts) a fly crossed the infrared beam running through the middle of each activity tube per 24-hour period. Activity data from the respectively aged flies were averaged and plotted as counts/24 hours.

#### Drug treatments

All drugs were purchased from Sigma-Aldrich (St. Louis, MO). For luciferase reporter assays, reserpine (300 μM final concentration) was diluted in dimethylsulfoxide (DMSO) and amphetamine (10 mM final concentration) was diluted in distilled water. Drugs and respective vehicle controls were mixed with molten cornmeal-molasses media and subsequently poured into standard fly vials.

#### Luciferase reporter assay

Adult DA neuron dVGLUT luciferase reporter flies were transferred into vials either with drug- or vehicle-treated food at 13 days post-eclosion and dissected 24 hours later or dissected at varying ages to analyze age-related changes in luciferase expression. Fly brains were rapidly dissected from decapitated flies with removal of associated cuticle and connective tissues with each sample consisting of 5 pooled fly brains. Luciferase assays were performed on samples using the Dual-Luciferase Reporter Assay System (Promega, Madison, WI). Brains were placed in 200 μL of 1x passive lysis buffer and homogenized with a pestle. Samples were then shaken at room temperature (45 min, 1200 rpm) and stored at -80°C until the assay was performed (within one month). Once thawed, respective sample lysates were vortexed and appropriately diluted in distilled water to ensure that luminescent signal was not oversaturated. 10μL of lysate from each sample was then added into a clear F-bottom 96-well plate in triplicate followed by addition of 100μL of Luciferase Assay Reagent II to each well in the dark. Wells were immediately placed in a Pherastar FSX plate reader (BMG Labtech, Ortenberg, Germany), focal height adjusted, and firefly luciferase-mediated luminescence was read. Firefly luciferase reactions were terminated with addition of 100 μL of Stop & Glo reagent which also provided an internal control via the *Renilla* luciferase present in the buffer as described earlier^29^. Triplicates for each sample were averaged, and results were reported as a luminescence ratio of firefly luciferase luminescence divided by *Renilla* luciferase luminescence (LUM ratio). Unless specified, luciferase assays used both male and female flies.

#### Multiplex fluorescent in situ hybridization

Multiplex fluorescent *in situ* hybridization was performed to measure mRNA expression as previously described^30^, with adaptations for labeling of adult central fly brain. Male w^1118^ fly brains were dissected 14 days post-eclosion in ice-cold Schneider’s insect medium and fixed in 2% paraformaldehyde in PBS. Following wash in PBS with 0.5% Triton X-100, brains were dehydrated in a graded series of ethanol dilutions and shaken in 100% ethanol (4°C, overnight). Brains were subsequently rehydrated with a series of ethanol solutions and incubated with 5% acetic acid to enhance probe penetration. Following PBS washing, brains were fixed again in 2% paraformaldehyde in PBS. After washing in PBST (PBS with 0.5% Triton X-100), brains were incubated with 1% NaBH_4_ at 4°C. After a final PBST wash, brains were transferred onto Superfrost Plus slides (Thermo Fisher Scientific) and air-dried. Multiplex fluorescent *in situ* hybridization via RNAscope was then performed according to manufacturer’s instructions (Advanced Cell Diagnostics, Hayward, CA) to detect mRNA expression of TH (*ple* gene, Cat. No. 536401-C2) and dVGLUT (*vglut* gene, Cat. No. 424011). Briefly, tissue sections were protease-treated, and then probes were hybridized to their target mRNAs (2 hours, 40°C). Sections were exposed to a series of incubations that amplified the target probes, and then counterstained with DAPI. dVGLUT and TH mRNAs were detected with Alexa 488 and Atto 550, respectively.

#### Confocal microscopy of Drosophila brain fluorescent in situ hybridization

Images were acquired with an Olympus IX81 inverted microscope equipped with an Olympus spinning disk confocal unit (Olympus, Center Valley, PA), a Hamamatsu EM-CCD digital camera (Hamamatsu, Bridgewater, NJ, USA) and a high-precision BioPrecision2 XYZ motorized stage with linear XYZ encoders (Ludl Electronic Products Ltd, Hawthorne, NJ) using a 60x 1.4 NA SC oil immersion objective. 3D image stacks (2048×2048 pixels, 0.2 μm z-steps) of the first 3 μm from the surface of the tissue thickness were taken to ensure full penetrance of probes. Image sites were systematically and randomly selected across the extent of each fly brain using a grid of 100 μm^2^ frames spaced by 200 μm. Image collection was controlled by Slidebook 6.0 (Intelligent Imaging Innovations, Inc., Denver, CO). The z-stacks were collected using optimal exposure settings (*i*.*e*., those that yielded the greatest dynamic range with no saturated pixels), with differences in exposures normalized during image processing.

#### Image analysis of mRNA expression

Imaging data was initially analyzed via Slidebook (Intelligent Imaging Innovations, Inc.) and Matlab (MathWorks, Natick, MA) software. First, a Gaussian channel was made for each channel for quantification of mRNA expression by calculating a difference of Gaussians using sigma values of 0.7 and 2. Then, an average projection of each 3D image stack was created by averaging intensity values within each Gaussian channel to assemble a 2D image of DAPI-stained cells and TH and VGLUT2 mRNA transcripts.

#### DAPI-stained cell and mRNA quantification

To quantify mRNA expression within the TH and dVGLUT channels, 2D projection images were separated into quantitative TIFF files of each individual Gaussian channel and transferred to the HALO image analysis platform equipped with a fluorescent *in situ* hybridization add-on (Version 1.7, Indica Labs, Albuquerque, NM). DAPI-stained cell nuclei and fluorescent grains representing mRNA transcripts from the VGLUT2 and TH channels were quantified via HALO software using the following parameters for inclusion in our counts based on the following thresholding criteria: any object 1-10 μm^2^ for DAPI and 0.03-0.15 μm^2^ for TH and dVGLUT grains. We quantified the respective expression levels of TH and dVGLUT mRNA in positive cells as described earlier^31^, with some modifications. To determine the minimum number of mRNA grains associated with a DAPI-stained nucleus considered positive, we tested different thresholds (TH: 50-, 100-, and 200-times; dVGLUT: 10-, 25-, and 50-times) of the number of mRNA grains above background levels (*i*.*e*., the number of grains expressed in a typical cell volume). Since there were no significant differences in relative TH^+^/dVGLUT^+^ cell densities between the thresholds (unpaired t-tests; all P>0.11), minimum thresholds of 100-times the background expression level for TH and 25-times the background expression level for dVGLUT were selected as the thresholds for quantifying positive cells. All mRNA grains within 1 μm of the nucleus edge were considered to belong to the respective cell, and this 1 μm border was reduced in HALO whenever necessary to prevent overlap between neighboring cells. TH and dVGLUT cell expression were quantified together and reported as density of TH^+^/dVGLUT^+^ cells.

#### Ex vivo multiphoton Drosophila brain imaging and analysis

Isolated *ex vivo* whole adult fly brain preparations were obtained by rapid brain removal and microdissection in adult hemolymph-like saline (AHL, in mM: 108 NaCl, 5 KCl, 2 CaCl_2_, 8.2 MgCl_2_, 1 NaH_2_PO_4_, 10 sucrose, 5 trehalose, 5 HEPES, 4 NaHCO_3_; pH 7.4, 265 mOsm) followed by pinning onto slabs of thin Sylgard with fine tungsten wire as described previously^10,19^. Under continuous AHL perfusion, the whole brain preparations were imaged on a Bergamo II resonant-scanning 2-photon microscope (ThorLabs Inc., Newton, NJ) using a SemiApo 20x (1.0 NA) water-immersion objective lens (Olympus, Tokyo, Japan). The illumination source was an Insight X3 IR laser (Spectra-Physics, Newport, Irvine, CA) mode-locked at 920 nm. Typically, <50mW mean power was delivered at the sample. Fluorescence emission was collected using a 550/50 nm full-width half-maximum (FWHM) bandpass emission filter for GFP (λ_ex_=920 nm). Z-stacks through the depth of the entire adult central brain were acquired in 2 μm steps using ThorImage 4.0 software (Thorlabs). Collected images had a voxel size of 0.64 (x) × 0.64 (y) × 2.00 (z) μm with 8x frame averaging. GFP-labeled cell bodies in the adult central fly brain were counted throughout all z-stacks from brains of both male and female flies± TH-driven dVGLUT RNAi at 2, 14 and 60 days post-eclosion. We focused on well-defined DA neuron clusters including those within the protocerebral anterior lateral (PAL), protocerebral anterior medial (PAM), and suboesophageal ganglion (SOG) regions^32-34^. For all cell body counts, comparable results were independently obtained from n≥3 blinded experimenters. Z-projections of the respective image stacks were generated for representative images using the Fiji/ImageJ image-processing package (National Institutes of Health, Bethesda, MD).

### Human Brain Experiments

#### Postmortem human subjects

Human postmortem brain specimens (N = 2 male and 2 female subjects) were obtained through the University of Pittsburgh Brain Tissue Donation Program. Brain tissue was obtained following consent from next-of-kin during autopsies conducted at the Allegheny County (Pittsburgh, PA) or Davidson County (Nashville, TN) Office of the Medical Examiner. An independent committee of experienced research clinicians confirmed the absence of lifetime psychiatric and neurologic diagnoses for all subjects on the basis of medical and neuropathological records and structured diagnostic interviews conducted with family members of the deceased^35,36^. Male and female subjects did not differ in mean age, postmortem interval, RNA integrity number, or brain pH (unpaired t-tests, all P>0.18; Extended Table 1). All procedures were approved by the University of Pittsburgh’s Committee for Oversight of Research and Clinical Training Involving Decedents and Institutional Review Board for Biomedical Research.

#### Fluorescent in situ hybridization

*In situ* hybridization probes for multiplex fluorescent *in situ* hybridization in human brain samples were designed by Advanced Cell Diagnostics to detect mRNAs encoding TH (*TH* gene, Cat. No. 441651-C2) and VGLUT2 (*SLC17A6* gene, Cat. No. 415671). Right hemisphere midbrain containing the VTA and SNc were cut at a 20 μm thickness using a cryostat, mounted onto Superfrost Plus slides (Thermo Fisher Scientific) and stored at -80°C until processing. Tissue sections (2 per subject) were processed within a month of sectioning, and one section from each subject was processed on the same day to reduce batch effects. Multiplex fluorescent *in situ* hybridization via RNAscope was performed according to manufacturer’s instructions (Advanced Cell Diagnostics). Briefly, tissue sections were fixed for 15 min in ice-cold 4% paraformaldehyde, incubated in a protease treatment, and then the probes were hybridized to their target mRNAs (2 hours, 40°C). The sections were exposed to a series of incubations that amplified the target probes, and then counterstained with DAPI. VGLUT2 and TH mRNAs were detected with Alexa 488 and Atto 550, respectively.

#### Confocal microscopy imaging of human brain sections processed for fluorescent in situ hybridization

Images were acquired with an Olympus IX81 inverted microscope equipped with an Olympus spinning disk confocal unit, a Hamamatsu EM-CCD digital camera and a high-precision BioPrecision2 XYZ motorized stage with linear XYZ encoders (Ludl Electronic Products Ltd) using a 60x 1.4 NA SC oil immersion objective. 3D image stacks (2048×2048 pixels; 0.2 μm z-steps) of 100% of the tissue thickness were taken in the VTA and SNc spanning the entire medial-lateral and dorsal-ventral axes. Image sites were systematically and randomly selected using a grid of 100 μm^2^ frames spaced by 350 μm. Image collection was controlled by Slidebook 6.0 (Intelligent Imaging Innovations, Inc.). The z-stacks were collected using optimal exposure settings (*i*.*e*., those that yielded the greatest dynamic range with no saturated pixels), with differences in exposures normalized during image processing. Lipofuscin, an intracellular lysosomal protein that accumulates with age^37,38^, is a major source of native fluorescence across the visible spectrum in human postmortem tissue; however, we effectively exclude lipofuscin signal from our RNAscope probe-specific signals in human samples using a previously described approach^39-44^. Specifically, we imaged lipofuscin using a fourth visible channel (excitation/emission: 405 nm/647 nm) and masked the lipofuscin signal using an optimal threshold value.

#### Image analysis of mRNA expression

Imaging data was initially analyzed via Slidebook (Intelligent Imaging Innovations, Inc.) and Matlab software (MathWorks). A Gaussian channel was made for each channel by calculating a difference of Gaussians using sigma values of 0.7 and 2. Average projections of each 3D image stack were subsequently created by averaging intensity values within each Gaussian channel to assemble a 2D image of DAPI-stained cells and TH and VGLUT2 mRNA transcripts. Objects from the other channels that overlapped with lipofuscin were eliminated from analyses by subtracting the lipofuscin Gaussian channel from the other channels.

#### Quantification of mRNA expression

To quantify mRNA expression within the TH and VGLUT2 channels, 2D projection images were separated into quantitative TIFF files of each individual Gaussian channel and transferred to the HALO image analysis platform equipped with a fluorescent *in situ* hybridization add-on (Version 1.7, Indica Labs). DAPI-stained cell nuclei and fluorescent grains representing mRNA transcripts from the VGLUT2 and TH channels were quantified using the following inclusion parameters based on our thresholding: any object 40-500 μm^2^ for DAPI and 0.1-0.5 μm^2^ for TH and VGLUT2. Positive cell expression levels were determined as previously described^31^. To determine the minimum number of mRNA grains associated with a DAPI-stained nucleus considered positive, we tested different thresholds of 1.5-, 3- and 5-times the number of mRNA grains above background levels. Because there were no significant differences in relative TH^+^/VGLUT2^+^ cell densities between the thresholds (unpaired t-tests; all P>0.36), a minimum threshold of 3-times the background expression level was chosen as the threshold for quantifying positive cells. All mRNA grains within 5 μm of the nucleus edge were considered as belonging to the respective cell, and this 5 μm border was reduced in HALO whenever necessary to prevent overlap between neighboring cells. TH and VGLUT2 mRNA expression in neurons within the VTA and SNc were quantified together and reported as density of TH^+^/VGLUT2^+^ cells.

### Rat Brain Experiments

#### Animals

Adult (10 month) male and female Lewis rats (Envigo, Indianapolis, IN) were maintained under standard temperature control conditions with a 12:12-hour light:dark cycle; conventional diet and water were available *ad libitum*. All experiments involving animal treatment and euthanasia were approved by the University of Pittsburgh Institutional Animal Care and Use Committee, which is fully accredited by the Association for Assessment and Accreditation of Laboratory Animal Care (AAALAC) International. Animals were cared for in accordance with all appropriate animal care guidelines according to the National Institutes of Health Animal Care and Use Program as well as the Animal Research: Reporting of *In Vivo* Experiments (ARRIVE) guidelines for reporting animal research. All efforts were made to ameliorate animal suffering.

#### Immunohistochemistry and image analysis

Rats were euthanized using pentobarbital, followed by transcardial perfusion and 4% paraformaldehyde fixation. Brains were fixed in paraformaldehyde for 24-hours and transferred to 30% sucrose at 4°C until sectioning. Nigral sections (35μm) were sliced on a freezing microtome and maintained by free-floating in cryoprotectant at -20°C until immunohistochemical labeling. Sections were labeled for TH (1:2000, AB1542, EMD Millipore, Burlington, MA) and VGLUT2 (1:500, 135403, Synaptic Systems, Goettingen, Germany), then mounted onto glass slides for imaging using a “primary antibody delete” (secondary antibody only) stained section to subtract background fluorescence as described earlier^45^. Images were acquired using an Olympus BX61 microscope and Fluoview 1000 software (Nikon, Melville, NY). Quantitative fluorescence measurements were monitored using standard operating imaging parameters to ensure the absence of saturated pixels during image acquisition. For quantitative comparisons, all imaging parameters (*e*.*g*., laser power, exposure, and pinhole) were held constant across specimens. At least 6 images were analyzed per animal with analysis performed using Nikon NIS-Elements Advanced Research software (Version 4.5; Nikon, Melville, NY). Results are reported as a count of VGLUT2 puncta within TH^+^ cells (# objects/TH^+^ Cell).

### Mouse Brain Experiments

#### Animals

We used male mice (8-12 weeks old) expressing Cre recombinase under the control of the DA transporter (DAT^cre^) (*Slc6a3*^*IRESCre*^, Jackson stock 006660, The Jackson Laboratory, Bar Harbor, ME) which were bred on a C57BL/6J genetic background and backcrossed to C57BL/6J for 12 generations. Mice were group-housed, and maintained on a 12:12-hour light:dark cycle with food and water available *ad libitum*. All mice were used in accordance with protocols approved by the University of California, San Diego (UCSD) Institutional Animal Care and Use Committee and cared for in accordance with both ARRIVE and NIH guidelines for laboratory animal care and safety.

#### Viruses

We used serotype 1, replication-incompetent, Adeno-associated viruses (AAV) to drive expression of VGLUT2 under the control of the EF1α promoter: AAV1-EF1α-DIO-VGLUT2 (referred to as AAV-VGLUT2 in results) as described previously^13^. The virus was packaged at the Salk GT3 vector core (La Jolla, CA). We also used serotype 5 AAV5-EF1α-DIO-VGLUT2-T2A-eGFP (referred to as AAV-VGLUT2-eGFP in results) as reported earlier^14^; this virus was packaged by Vigene Biosciences (Rockville, MD).

#### Viral injections

Mice were anaesthetized with isoflurane (2-5%) and placed into a stereotaxic frame (David Kopf Instruments, Tujunga, CA). 300 nl of the following viruses were used: AAV-VGLUT2 (3.3×10^12^ genome copies per ml [gc/ml]), AAV-VGLUT2-eGFP (5×10^13^ gc/ml; concentrated titer), or PBS-diluted AAV-VGLUT2-eGFP (2×10^13^ gc/ml; originally reported titer^14^). The respective viruses were microinfused into the left SNc (−3.4 anterior-posterior, -1.25 medial-lateral, -4.25 dorsal-ventral; in millimeters from bregma) using custom-made 30G stainless steel injectors at a speed of 100nl/min.

#### Immunohistochemistry

Mice were deeply anaesthetized with pentobarbital (200 mg/kg i.p.; Virbac Corp., Westlake, TX) and transcardially perfused with 10–20ml of phosphate-buffered saline (PBS) 3 weeks after viral injection. This was followed by perfusion with 60-70ml of 4% paraformaldehyde (PFA) at a rate of 6 ml/min. Brains were extracted, post-fixed in 4% PFA at 4°C overnight, and cryoprotected in 30% sucrose in PBS for 48–72 hours at 4°C. Brains were snap-frozen in chilled isopentane and stored at -80°C. Sections (30μm) were sectioned using a cryostat (CM3050S, Leica, Wetzlar, Germany) and collected in PBS containing 0.01% sodium azide.

For fluorescent immunostaining, brain sections were blocked with 5% normal donkey serum in PBS containing 0.3% Triton X-100 (blocking buffer) (1-hour, room temperature). Sections were then incubated with one or more of the following primary antibodies (rabbit anti-TH, 1:2000, AB152, EMD Millipore; guinea pig anti-VGLUT2, AB2251, EMD Millipore; chicken anti-GFP, A10262, Invitrogen, Carlsbad, CA) in blocking buffer (overnight, 4°C). Sections were rinsed 3×15 min with PBS and incubated in appropriate secondary antibodies (Jackson ImmunoResearch, West Grove, PA) conjugated to Alexa 488, Alexa 594 or Alexa 647 fluorescent dyes (5 μg/ml) (2-hours, room temperature). Sections were washed 3×15 min with PBS, mounted onto glass slides and coverslipped with Fluoromount-G mounting medium (Southern Biotech, Birmingham, AL) supplemented with DAPI stain (0.5 µg/ml, Roche, Basel, Switzerland). Images were acquired using a Zeiss AxioObserver epifluorescence microscope (Oberkochen, Germany).

For TH-DAB staining, sections were quenched in 3% H_2_O_2_ in PBS (30 min, room temperature) before blocking (1-hour, room temperature). Sections were incubated with rabbit anti-TH primary antibody (1:2000; AB152, Millipore) in blocking buffer (overnight, 4°C). The following day, sections were washed 3×15 min with PBS and incubated with a donkey anti-rabbit biotinylated secondary antibody (Jackson ImmunoResearch) at 1:500 in blocking buffer (2-hours, room temperature).

Sections were again washed 3×15 min with PBS and incubated in avidin-biotin complex solution (Vectastain Elite ABC kit, Vector Laboratories, Burlingame, CA) (2-hours, room temperature) before 2 additional PBS washes (10 minutes). Sections were incubated in DAB solution (0.4mg/ml 3,3-diaminobenzidine-HCl, 0.005% H_2_O_2_ in PBS) (3 min, room temperature). Sections were rinsed twice in PBS before mounting onto glass slides and drying overnight. Sections were then dehydrated through increasing concentrations of ethanol and isopropanol, cleared with CitriSolv (Thermo Fisher Scientific, Waltham, MA), and cover-slipped using DPX mounting medium (Sigma-Aldrich, St. Louis, MO).

#### Unbiased stereology

Stereological sampling was performed using the Stereo Investigator (SI) software (MBF Bioscience, Williston, VT). Counting frames (100×100µm) were randomly placed on a counting grid (200×200 µm) and sampled using a 7-µm optical dissector with guard zones of 10% of the total slice thickness on each site (~2 µm). The boundaries of SNc were outlined under magnification (4x objective). Cells were counted with a 20x objective using a Zeiss AxioImager microscope (Carl Zeiss Microscopy, White Plains, NY). A DAergic neuron was defined as an in-focus TH-DAB-immunoreactive (TH-IR) cell body with a TH-negative nucleus within the counting frame. Every sixth section was processed for TH-IR, resulting in 6-7 sections containing SNc sampled per mouse and every section was counted. The number of neurons in SNc was estimated using the optical fractionator method, which is unaffected by changes in the volume of reference of the structure sampled.

#### TH and VGLUT2 densitometry

Images (TH-DAB stained) were acquired using a Zeiss AxioObserver microscope (Carl Zeiss Microscopy) under brightfield illumination. Four striatal sections per animal were analyzed using ImageJ software (National Institutes of Health). Regions of interest in the dorsal striatum were delineated, pixel densities were estimated and intensities over the four sections were averaged.

### Statistics

Drug- and sex-mediated changes in luciferase reporter expression in flies, sex differences in rat VGLUT2 expression and hemisphere comparisons after AAV-mediated striatal VGLUT2 overexpression were analyzed via unpaired Student’s t-test. Age-related changes in luciferase reporter luminescence and striatal VGLUT2 comparisons between mouse viral vectors were analyzed via one-way analysis of variance (ANOVA). *Drosophila* locomotion and DA neuron numbers in the absence of RNAi-mediated dVGLUT knockdown were analyzed using two-way ANOVA, with sex and age as between-subjects factors. In dVGLUT RNAi flies, locomotion and DA neuron number were analyzed using three-way ANOVA with sex, age and dVGLUT RNAi as between-subjects factors. DA neuron survival and density of striatal DAergic projections after AAV injection were analyzed using repeated measures two-way ANOVA with AAV preparation as a between-subjects factor and hemisphere relative to injection as a within-subjects factor. Pearson correlation coefficients were calculated to correlate AAV-mediated VGLUT2 overexpression to TH^+^ DA neuron survival and striatal TH^+^ terminal density. Significant effects were followed up with Bonferroni post-hoc comparisons. In all analyses, statistical significance was defined as P<0.05. All statistical functions were completed using GraphPad Prism software (version 8.2, GraphPad Software, San Diego, CA).

## Supplemental Information

Extended data includes five figures and a table.

## Acknowledgements

We thank Hui Shen, Timothy Crawley, Emily W. George and Alisa Pugacheva for assistance. This work was supported by awards from the National Institutes of Health R21AG068607 (Z.F.), K08DA031241 (Z.F.), K99AG059834 (T.S.), K99ES029986 (B.R.D.M.), T32NS07433 (S.A.B.), T32GM008208 (E.I.O.L.), the Louis V. Gerstner, Jr., Scholars Program (Z.F.), Leon Levy Foundation (Z.F.), the John F. and Nancy A. Emmerling Fund of The Pittsburgh Foundation (Z.F.), Austrian Science Fund/FWF Schrödinger fellowship J3656-B24 (T.S.), and the American Parkinson Disease Association Center for Advanced Research at the University of Pittsburgh (J.T.G.).

## Author Contributions

S.A.B. and Z.F. conceived the project. S.A.B., D.A., M.V., S.A.B., V.C.C., S.A.R., E.I.O.L., E.G.N., K.J.F., M.J.P., R.W.L., B.D.M., C.E.J.C., and Z.F. performed *Drosophila* genetics, imaging, and behavioral experiments with accompanying data analysis. B.R.D.M., S.A.B., and J.T.G. performed rat experiments. T.S., S.A.B., A.B., and T.S.H. performed all mouse experiments. S.A.B. and Z.F. wrote the manuscript with contributions from co-authors.

## Competing Interests

The authors report no competing interests.

## Notes

### Competing Interest Statement

The authors have declared no competing interest.

## References

1 Hartmann, A. Postmortem studies in Parkinson’s disease. Dialogues Clin Neurosci 6, 281–293 (2004).

2 Reeve, A., Simcox, E. & Turnbull, D. Ageing and Parkinson’s disease: why is advancing age the biggest risk factor? Ageing Res Rev 14, 19–30, doi:10.1016/j.arr.2014.01.004 (2014).

3 Hirsch, E., Graybiel, A. M. & Agid, Y. A. Melanized dopaminergic neurons are differentially susceptible to degeneration in Parkinson’s disease. Nature 334, 345–348, doi:10.1038/334345a0 (1988).

4 Brichta, L. & Greengard, p. Molecular determinants of selective dopaminergic vulnerability in Parkinson’s disease: an update. Frontiers in neuroanatomy 8, 152, doi:10.3389/fnana.2014.00152 (2014).

5 Giguere, N., Burke Nanni, S. & Trudeau, L. E. On Cell Loss and Selective Vulnerability of Neuronal Populations in Parkinson’s Disease. Frontiers in neurology 9, 455, doi:10.3389/fneur.2018.00455 (2018).

6 Surmeier, D. J. Determinants of dopaminergic neuron loss in Parkinson’s disease. The FEBS journal 285, 3657–3668, doi:10.1111/febs.14607 (2018).

7 Sulzer, D. & Surmeier, D. J. Neuronal vulnerability, pathogenesis, and Parkinson’s disease. Movement disorders : official journal of the Movement Disorder Society 28, 715–724, doi:10.1002/mds.25187 (2013).

8 Trudeau, L. E. & El Mestikawy, S. Glutamate Cotransmission in Cholinergic, GABAergic and Monoamine Systems: Contrasts and Commonalities. Frontiers in neural circuits 12, 113, doi:10.3389/fncir.2018.00113 (2018).

9 Mingote, S., Amsellem, A., Kempf, A., Rayport, S. & Chuhma, N. Dopamine-glutamate neuron projections to the nucleus accumbens medial shell and behavioral switching. Neurochemistry international 129, 104482, doi:10.1016/j.neuint.2019.104482 (2019).

10 Aguilar, J. I. et al. Neuronal Depolarization Drives Increased Dopamine Synaptic Vesicle Loading via VGLUT. Neuron 95, 1074–1088 e1077, doi:10.1016/j.neuron.2017.07.038 (2017).

11 Hnasko, T. S. et al. Vesicular glutamate transport promotes dopamine storage and glutamate corelease in vivo. Neuron 65, 643–656, doi:10.1016/j.neuron.2010.02.012 (2010).

12 Kouwenhoven, W. M. et al. VGluT2 Expression in Dopamine Neurons Contributes to Postlesional Striatal Reinnervation. J Neurosci 40, 8262–8275, doi:10.1523/jneurosci.0823-20.2020 (2020).

13 Steinkellner, T. et al. Role for VGLUT2 in selective vulnerability of midbrain dopamine neurons. J Clin Invest 128, 774–788, doi:10.1172/jci95795 (2018).

14 Shen, H. et al. Genetic deletion of vesicular glutamate transporter in dopamine neurons increases vulnerability to MPTP-induced neurotoxicity in mice. Proc Natl Acad Sci U S A 115, E11532–e11541, doi:10.1073/pnas.1800886115 (2018).

15 Yamamoto, S. & Seto, E. S. Dopamine dynamics and signaling in Drosophila: an overview of genes, drugs and behavioral paradigms. Experimental animals / Japanese Association for Laboratory Animal Science 63, 107–119 (2014).

16 Bainton, R. J. et al. Dopamine modulates acute responses to cocaine, nicotine and ethanol in Drosophila. Curr Biol 10, 187–194 (2000).

17 Mingote, S. et al. Dopamine neuron dependent behaviors mediated by glutamate cotransmission. eLife 6, doi:10.7554/eLife.27566 (2017).

18 Yamaguchi, T., Sheen, W. & Morales, M. Glutamatergic neurons are present in the rat ventral tegmental area. The European journal of neuroscience 25, 106–118, doi:10.1111/j.1460-9568.2006.05263.x (2007).

19 Freyberg, Z. et al. Mechanisms of amphetamine action illuminated through optical monitoring of dopamine synaptic vesicles in Drosophila brain. Nature communications 7, 10652, doi:10.1038/ncomms10652 (2016).

20 Neckameyer, W. S., Woodrome, S., Holt, B. & Mayer, A. Dopamine and senescence in Drosophila melanogaster. Neurobiology of aging 21, 145–152, doi:10.1016/s0197-4580(99)00109-8 (2000).

21 Friggi-Grelin, F. et al. Targeted gene expression in Drosophila dopaminergic cells using regulatory sequences from tyrosine hydroxylase. J Neurobiol 54, 618–627, doi:10.1002/neu.10185 [doi] (2003).

22 Berry, J. A., Cervantes-Sandoval, I., Chakraborty, M. & Davis, R. L. Sleep Facilitates Memory by Blocking Dopamine Neuron-Mediated Forgetting. Cell 161, 1656–1667, doi:10.1016/j.cell.2015.05.027 (2015).

23 Sherer, L. M. et al. Octopamine neuron dependent aggression requires dVGLUT from dual-transmitting neurons. PLoS Genet 16, e1008609, doi:10.1371/journal.pgen.1008609 (2020).

24 Diao, F. et al. Plug-and-play genetic access to drosophila cell types using exchangeable exon cassettes. Cell reports 10, 1410–1421, doi:10.1016/j.celrep.2015.01.059 (2015).

25 Yeh, E., Gustafson, K. & Boulianne, G. L. Green fluorescent protein as a vital marker and reporter of gene expression in Drosophila. Proc Natl Acad Sci U S A 92, 7036–7040 (1995).

26 Choi, B. J. et al. Miniature neurotransmission regulates Drosophila synaptic structural maturation. Neuron 82, 618–634, doi:10.1016/j.neuron.2014.03.012 (2014).

27 Nern, A., Pfeiffer, B. D., Svoboda, K. & Rubin, G. M. Multiple new site-specific recombinases for use in manipulating animal genomes. Proc Natl Acad Sci U S A 108, 14198–14203, doi:10.1073/pnas.1111704108 (2011).

28 Wang, J. W., Beck, E. S. & McCabe, B. D. A modular toolset for recombination transgenesis and neurogenetic analysis of Drosophila. PLoS One 7, e42102, doi:10.1371/journal.pone.0042102 (2012).

29 Hirose, F., Ohshima, N., Kwon, E. J., Yoshida, H. & Yamaguchi, M. Drosophila Mi-2 negatively regulates dDREF by inhibiting its DNA-binding activity. Molecular and cellular biology 22, 5182–5193, doi:10.1128/mcb.22.14.5182-5193.2002 (2002).

30 Long, X., Colonell, J., Wong, A. M., Singer, R. H. & Lionnet, T. Quantitative mRNA imaging throughout the entire Drosophila brain. Nat Methods 14, 703–706, doi:10.1038/nmeth.4309 (2017).

31 Rocco, B. R., Sweet, R. A., Lewis, D. A. & Fish, K. N. GABA-Synthesizing Enzymes in Calbindin and Calretinin Neurons in Monkey Prefrontal Cortex. Cereb Cortex 26, 2191–2204, doi:10.1093/cercor/bhv051 (2016).

32 Nassel, D. R. & Elekes, K. Aminergic neurons in the brain of blowflies and Drosophila: dopamine- and tyrosine hydroxylase-immunoreactive neurons and their relationship with putative histaminergic neurons. Cell Tissue Res 267, 147–167, doi:10.1007/bf00318701 (1992).

33 Mao, Z. & Davis, R. L. Eight different types of dopaminergic neurons innervate the Drosophila mushroom body neuropil: anatomical and physiological heterogeneity. Frontiers in neural circuits 3, 5, doi:10.3389/neuro.04.005.2009 (2009).

34 Marella, S., Mann, K. & Scott, K. Dopaminergic modulation of sucrose acceptance behavior in Drosophila. Neuron 73, 941–950, doi:10.1016/j.neuron.2011.12.032 [doi] S0896-6273(12)00082-7 [pii] (2012).

35 Glantz, L. A. & Lewis, D. A. Decreased dendritic spine density on prefrontal cortical pyramidal neurons in schizophrenia. Arch Gen Psychiatry 57, 65–73, doi:10.1001/archpsyc.57.1.65 (2000).

36 Glausier, J. R., Kelly, M. A., Salem, S., Chen, K. & Lewis, D. A. Proxy measures of premortem cognitive aptitude in postmortem subjects with schizophrenia. Psychol Med 50, 507–514, doi:10.1017/S0033291719000382 (2020).

37 Benavides, S. H., Monserrat, A. J., Farina, S. & Porta, E. A. Sequential histochemical studies of neuronal lipofuscin in human cerebral cortex from the first to the ninth decade of life. Arch Gerontol Geriatr 34, 219–231, doi:10.1016/s0167-4943(01)00223-0 (2002).

38 Porta, E. A., Berra, A., Monserrat, A. J. & Benavides, S. H. Differential lectin histochemical studies on lipofuscin (age-pigment) and on selected ceroid pigments. Arch Gerontol Geriatr 34, 193–203, doi:10.1016/s0167-4943(01)00224-2 (2002).

39 Sweet, R. A., Fish, K. N. & Lewis, D. A. Mapping Synaptic Pathology within Cerebral Cortical Circuits in Subjects with Schizophrenia. Front Hum Neurosci 4, 44, doi:10.3389/fnhum.2010.00044 (2010).

40 Curley, A. A. et al. Cortical deficits of glutamic acid decarboxylase 67 expression in schizophrenia: clinical, protein, and cell type-specific features. Am J Psychiatry 168, 921–929, doi:10.1176/appi.ajp.2011.11010052 (2011).

41 Glausier, J. R., Fish, K. N. & Lewis, D. A. Altered parvalbumin basket cell inputs in the dorsolateral prefrontal cortex of schizophrenia subjects. Mol Psychiatry 19, 30–36, doi:10.1038/mp.2013.152 (2014).

42 Rocco, B. R., Lewis, D. A. & Fish, K. N. Markedly Lower Glutamic Acid Decarboxylase 67 Protein Levels in a Subset of Boutons in Schizophrenia. Biol Psychiatry 79, 1006–1015, doi:10.1016/j.biopsych.2015.07.022 (2016).

43 Rocco, B. R., DeDionisio, A. M., Lewis, D. A. & Fish, K. N. Alterations in a Unique Class of Cortical Chandelier Cell Axon Cartridges in Schizophrenia. Biol Psychiatry 82, 40–48, doi:10.1016/j.biopsych.2016.09.018 (2017).

44 Fish, K. N., Rocco, B. R. & Lewis, D. A. Laminar Distribution of Subsets of GABAergic Axon Terminals in Human Prefrontal Cortex. Frontiers in neuroanatomy 12, 9, doi:10.3389/fnana.2018.00009 (2018).

45 De Miranda, B. R., Fazzari, M., Rocha, E. M., Castro, S. & Greenamyre, J. T. Sex Differences in Rotenone Sensitivity Reflect the Male-to-Female Ratio in Human Parkinson’s Disease Incidence. Toxicological sciences : an official journal of the Society of Toxicology 170, 133–143, doi:10.1093/toxsci/kfz082 (2019).

