## Supplemental Extended Data for "VGLUT modulates sex differences in dopamine neuron vulnerability to age-related neurodegeneration"

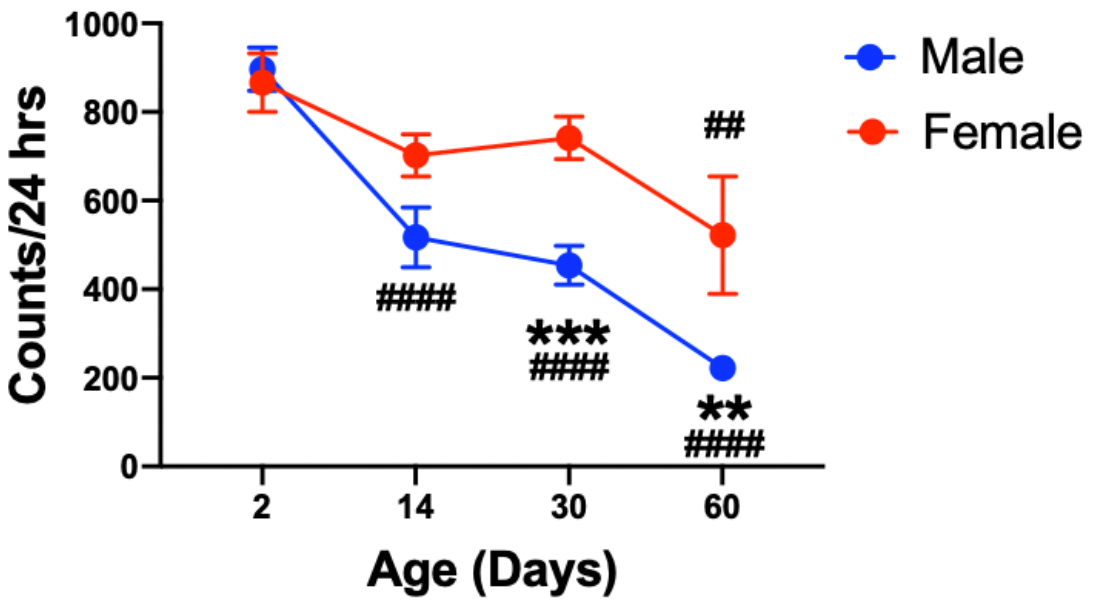


**Extended Data Fig. 1. Sex differences in age-dependent locomotor loss in *Drosophila*.** Adult male flies demonstrate progressive loss of 24-hour locomotion across aging compared to females who only show diminished locomotion at day 60 post-eclosion. Two-way ANOVA: Age: $F_{3,169}$=26.9, P<0.0001; Sex: $F_{1,169}$=20.7, P<0.0001; Age×Sex: $F_{3,169}$=4.1, P=0.0072. Results represented as mean±SEM; **P<0.01, ***P<0.001 compared to opposite sex, $^{\#\#}$P<0.01, $^{\#\#\#\#}$P<0.0001 compared to locomotion in 2-day-old flies by Bonferroni post-hoc comparison N=11-31 animals per group.

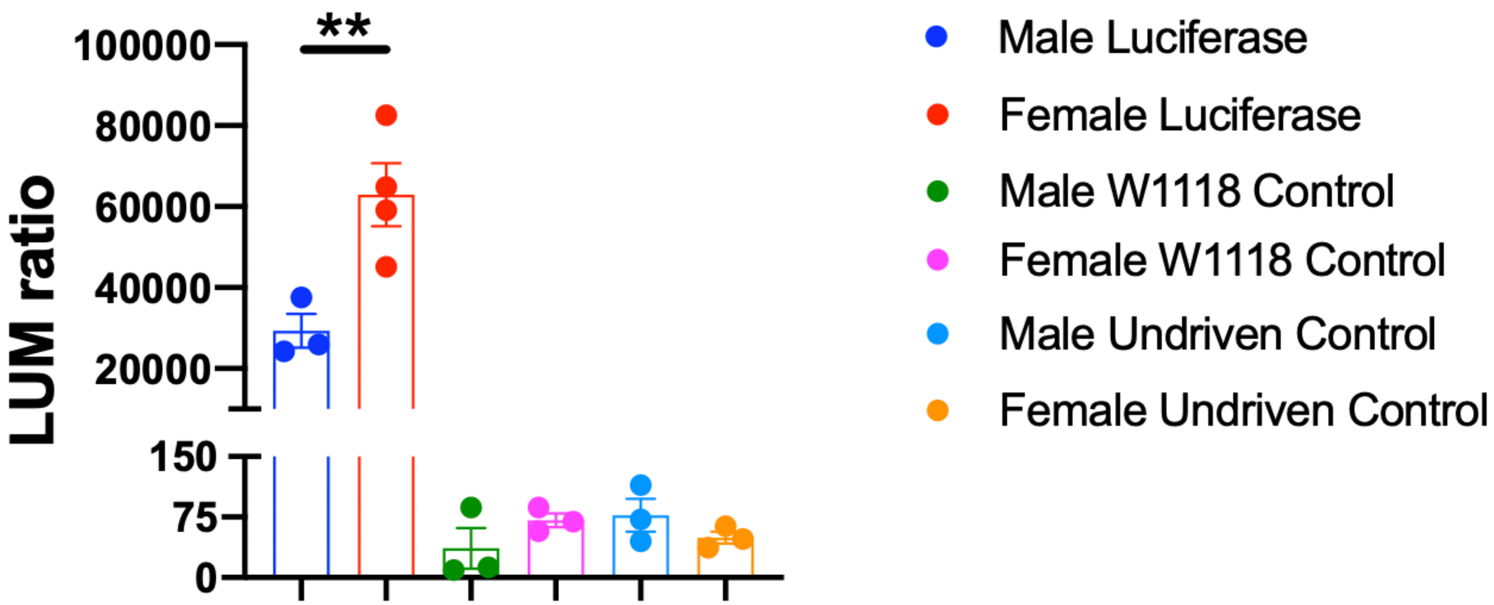


**Extended Data Fig. 2. Original intersectional DA neuron dVGLUT luciferase reporter luminescence and controls.** Female flies express 2-fold more dVGLUT in DA neurons compared to males. The TH-LexA and dVGLUT-GAL4-driven intersectional genetic reporter of DA neuron dVGLUT expression expresses 1200-fold more luminescence compared to the undriven (LexAop-B3R/+;UAS-B3RT.STOP.B3RT.Luciferase/+) or wild-type $w^{1118}$ controls of either sex. There are no significant differences in firefly:Renilla luminescence ratios between the controls (one-way ANOVA: $F_{3,8}$=1.2, P>0.05). Points represent individual animals. Results represented by mean±SEM; *P<0.05 by unpaired t-test, N=3-7 per group.

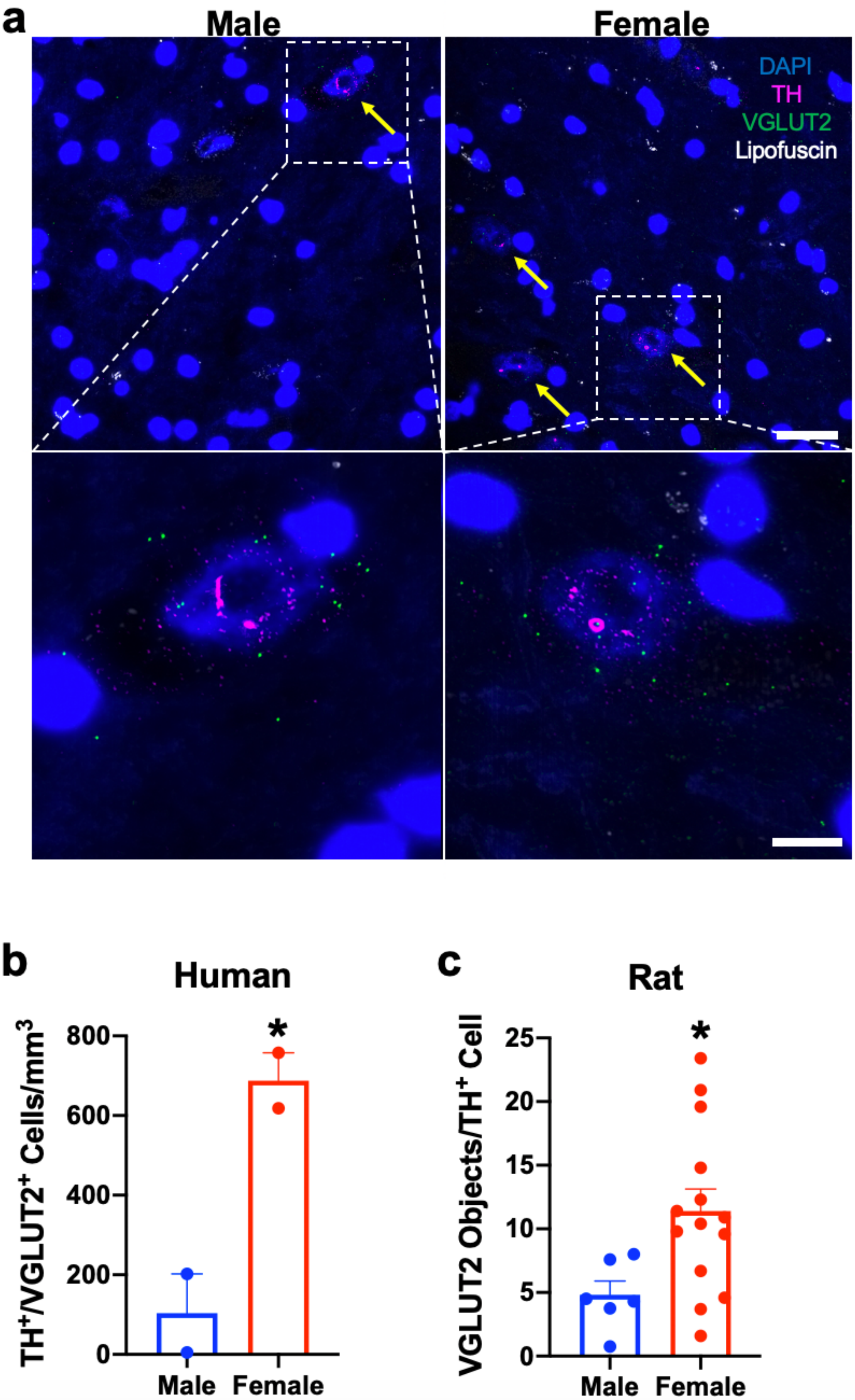
a
Male
Female
DAPI
TH
VGLUT2
Lipofuscin
b
Human
TH+/VGLUT2+ Cells/mm3
800
600
400
200
0
*
Male
Female
c
Rat
VGLUT2 Objects/TH+ Cell
25
20
15
10
5
0
*
Male
Female

**Extended Data Fig. 3. Sex differences in human and rat DA neuron VGLUT2 expression. (a)** Representative images of TH (magenta) and VGLUT2 (green) mRNA expression in postmortem human VTA/SNc of male and female subjects; yellow arrows denote $TH^+/VGLUT2^+$ neurons. Whole field image scale bar=25 μm, inset scale bar=10 μm. **(b)** Human $TH^+/VGLUT2^+$ neuron density is 6.6 times higher in female subjects compared to males (two-tailed unpaired t-test: $t_2$=4.8, P=0.040). N=2 subjects per group. **(c)** Rat $TH^+$ midbrain neurons show 2.4-fold elevated VGLUT2 expression in females compared to males (two-tailed unpaired t-test: $t_{18}$=2.4, P=0.029). N=6 males, 14 females. Results normalized to males. Points represent individual animals or subjects. Results represented as mean±SEM; *P<0.05 by unpaired t-test.

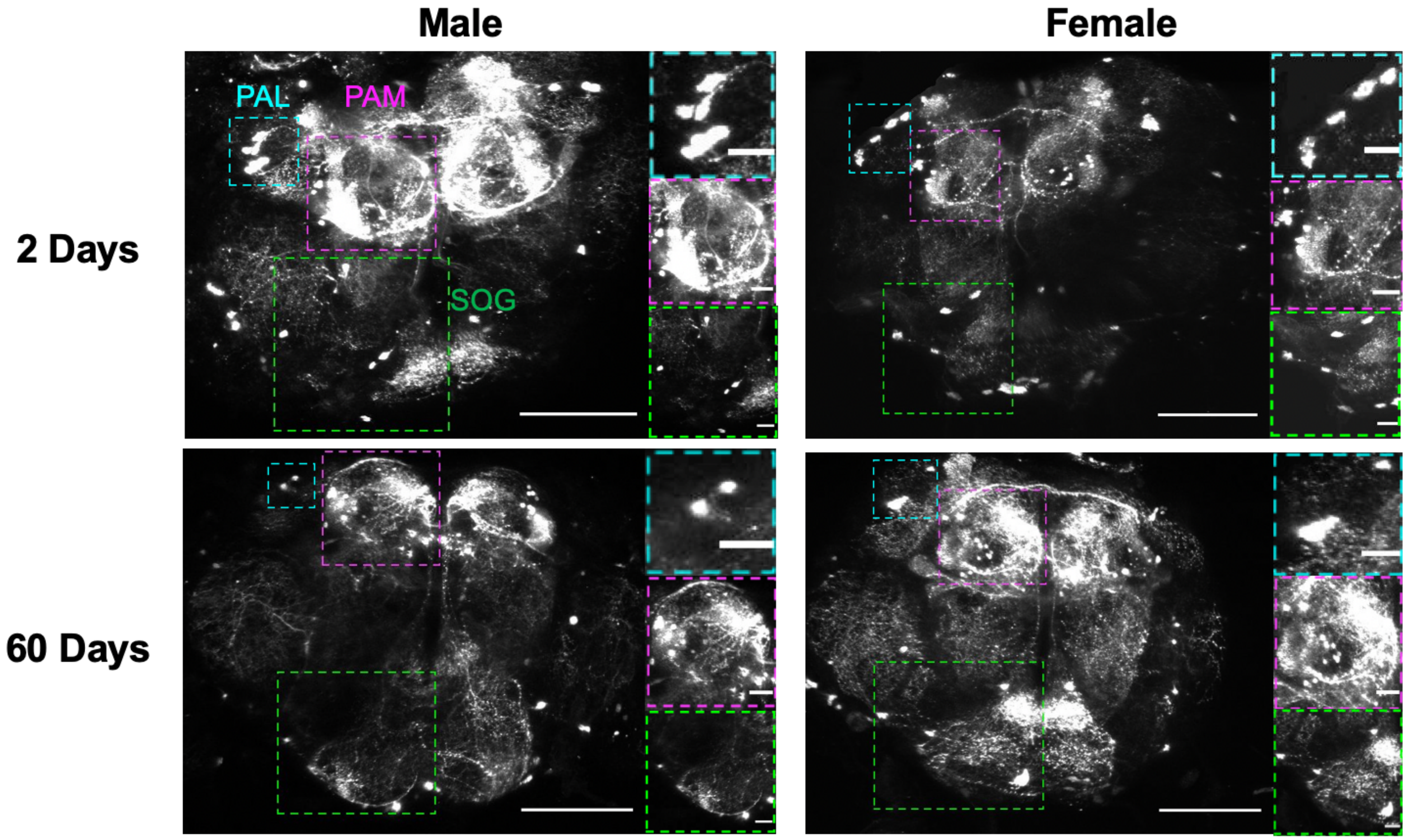


**Extended Data Fig. 4. Representative images of GFP-labeled DA neurons in whole intact adult male and female *Drosophila* brains.** Representative two-dimensional (2D) projection images of multiphoton microscopy of living whole fly brains of male and female adult flies (TH-GAL4/UAS-GFP) aged 2- versus 60-days post-eclosion. Insets highlight SOG (green box), PAL (white box), and PAM (magenta box) DA neuron clusters and show age-related loss of PAL and SOG DA neurons in males, but not females. Scale bars of 2D whole brain projections=100 μm, scale bars of insets=20 μm.

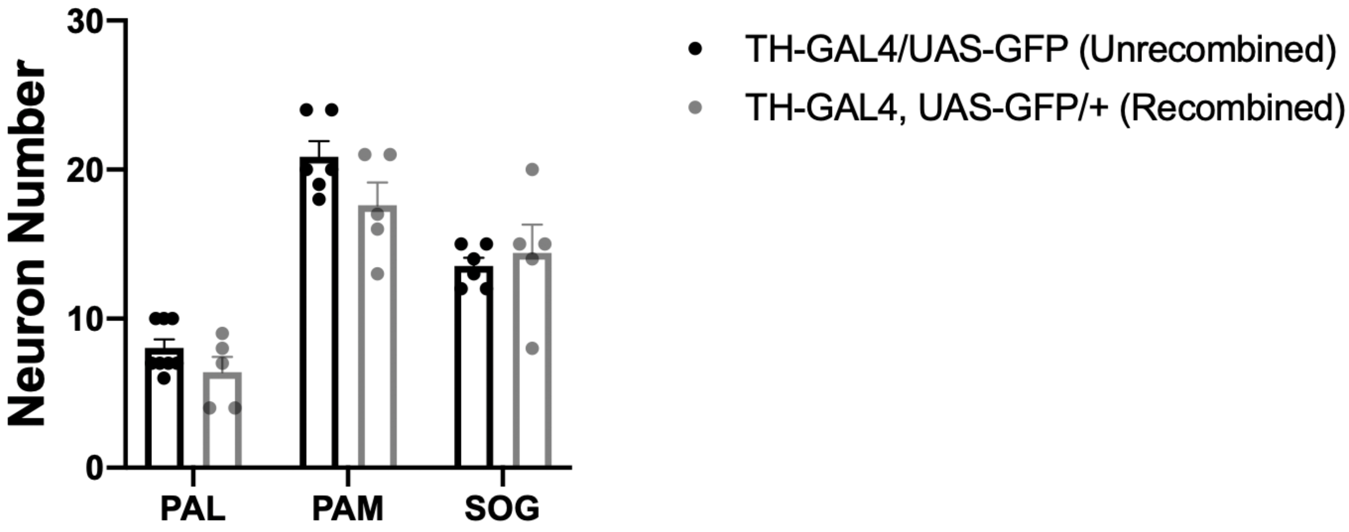


**Extended Data Fig. 5. Recombination of TH-GAL4 and UAS-GFP does not alter numbers of labeled DA neurons.** DA neurons were counted in DA neuron clusters in the PAL, PAM and SOG brain regions imaged by multiphoton microscopy of whole living *Drosophila* brains of 2-day-old adult males. DA neurons were labeled by combining the TH-GAL4 expression driver and UAS-GFP via genetic recombination (TH-GAL4, UAS-GFP/+; termed 'Recombined'). The DA neurons in brains of Recombined flies were compared to GFP-labeled DA neurons in unrecombined flies (TH-GAL4/UAS-GFP; termed 'Unrecombined'). There is no significant difference in $GFP^+$ DA neuron number in the PAL, PAM and SOG regions of Recombined versus Unrecombined brains (Bonferroni post-hoc test: P>0.05 for all regions). Points represent individual animals. Results represented as mean±SEM; N=4-8 brains per group.

| Subject | Sex | Age (years) | Postmortem interval (hours) | RNA integrity number | Brain pH |
|---|---|---|---|---|---|
| 1 | Male | 17 | 15.1 | 6.9 | 7.9 |
| 2 | Male | 22 | 20.1 | 6.9 | 7.8 |
| 3 | Female | 16 | 9.3 | 6.6 | 9.0 |
| 4 | Female | 21 | 23.9 | 6.8 | 8.2 |

**Extended Table 1. Characteristics of human subjects.** Male and female subjects were matched with no significant differences in mean age (males: 19.5±3.5 years, females: 18.5±3.5), postmortem interval (males: 17.6±3.5 hours, females: 16.6±10.3 hours), RNA integrity number (males: 7.9±0.1, females: 8.6±0.6) or brain pH (males: 6.9±0.01, females: 6.7±0.1); P>0.18 for all comparisons.
